# CuAAC stabilization of an NMR mixed labeled dimer

**DOI:** 10.1101/2021.05.24.445505

**Authors:** Paul J. Sapienza, Michelle M. Currie, Kelin Li, Jeffrey Aubé, Andrew L. Lee

## Abstract

Homo dimers are the most abundant type of enzyme in cells and as such, they represent the archetypal system for studying the remarkable phenomenon of allostery. In these systems, in which the allosteric features are manifest by the effect of the first binding event on the similar event at the second site, the most informative state is the asymmetric single bound (lig_1_) form, yet it tends to be elusive thermodynamically. Here we take significant steps towards obtaining milligram quantities of pure lig_1_ of the allosteric homodimer, chorismate mutase, in the form of a mixed isotopically labeled dimer stabilized by Cu^(I)-^catalyzed azide–alkyne cycloaddition (CuAAC) between the subunits. Below, we outline several critical steps required to generate high yields of both types of unnatural amino acid-containing proteins, and overcome multiple pitfalls intrinsic to CuAAC to obtain high yields of pure, fully intact, and active mixed labeled dimer. These data not only will make possible NMR-based investigations of allostery envisioned by us, but should also facilitate other structural applications where specific linkage of proteins is helpful.

## Introduction

Allostery is a remarkable phenomenon that is used ubiquitously in nature to achieve homeostasis. The motivation to understand allostery comes from a desire to untangle the native mechanisms of protein function, as well as the ultimate goal of designing allosteric proteins and allosteric drugs^1–4^. It is now clear that both structure and dynamics must be accounted for in order to describe allosteric transitions, and NMR is the single tool that is most ideally suited to provide both types of information^5^. Nature often uses oligomerization and symmetry in allosteric mechanisms, and because homodimers are the most common form of enzyme found in nature^6, 7^, we have chosen to study homodimeric systems by NMR to gain a deeper understanding of allostery. In the case of a symmetrical homodimer, where the allosteric feature is manifest by the effect of the first binding or catalytic event on the similar event on the second site, the most informative state is the asymmetric single bound (lig_1_) state. The “dream experiment” would be to bind ligand/substrate in one subunit while leaving the other subunit empty, and then observe how the binding of first ligand induces changes to either subunit, followed by similar observations corresponding to binding of the second ligand. Unfortunately, this most informative lig_1_ state, is also the most elusive to capture in highly abundant and/or purified form, because, except for cases of extreme negative cooperativity, the lig_1_ state is disfavored^8^. To perform NMR studies on the lig_1_ state, we pioneered a mixed labeled dimer (MLD) approach in which isotopically labeled dimeric thymidylate synthase was mixed with unlabeled and binding incompetent dimer having a different charge, and the MLD was purified using ion exchange chromatography^9^. We were able to perform some NMR experiments on the MLD because the kinetics of dimer reapportionment in thymidylate synthase was slow. However, even in this favorable case, return to the equilibrium manifold of dimeric states occurred sufficiently fast to prevent conducting more time consuming and informative spin relaxation experiments. We therefore sought a method to selectively covalently link MLD subunits and trap them in the desired state.

We chose the homodimeric enzyme chorismate mutase from *Saccharomyces cerevisiae* (Figure 1) to explore the linked MLD approach because of its rich allosteric features and the fact that CM subunits exchange rapidly in the absence of linkage. CM is a classically allosteric protein that falls in the biosynthesis of aromatic amino acids (Shikimate pathway), and it converts chorismic acid to prephenate^10^. Its activity is positively cooperative with respect to chorismate concentration (homotropic cooperativity), and the activity profile increases or decreases when bound to tryptophan or tyrosine, respectively (heterotropic cooperativity)^11, 12^. We weighed two approaches to covalently link CM subunits, either disulfide-based, or click chemistry together with unnatural amino acids (UAAs). Cystine disulfide bond linkage has the advantages of being reversible and not depending on UAA technology, but has limitations in that it is not biorthogonal, requiring the replacement of native Cys residues and/or omitting reducing agents, which can compromise function, replacing Cys residues prevents other NMR-based methods requiring thiol chemistry such as paramagnetic relaxation enhancement^13^, and finally, the short Cys side chain places severe constraints on the “reach” of linking groups. We chose the click-based approach, specifically the Cu^(I)^-catalyzed azide–alkyne cycloaddition (CuAAC) reaction, because it is proceeds rapidly in a diverse set of solution conditions, it is biorthogonal^14^, and there is a diverse set of UAAs with different geometries that may satisfy different structural demands on linkage^15^. There are drawbacks to the CuAAC approach, however. CuAAC requires incorporation of two different UAAs, one with an azide and the other containing an alkyne. This is straightforward in principle using amber suppression and engineered tRNA synthetases^16^. In practice, nearly all CuAAC applications in biological systems are successful on the basis of the need to only have one protein containing a single UAA. In such cases, the other click partner is supplied as a small molecule that is either synthesized or commercially available and therefore can be supplied in high concentration. Most of the time, complete reactivity of the macromolecule is not required to achieve the desired outcome, which is true from the early seminal modification of viral capsids^17^ to modern proteomic applications^18^. In structural protein-protein interaction uses, such as the NMR approach described here, high yield of proteins with *two different* UAAs is required, as is homogeneity of clicked species; homogeneity is difficult because Cu^I^-associated reaction oxygen species (ROS) often damage macromolecules during the reaction^19^.

**Figure 1.**
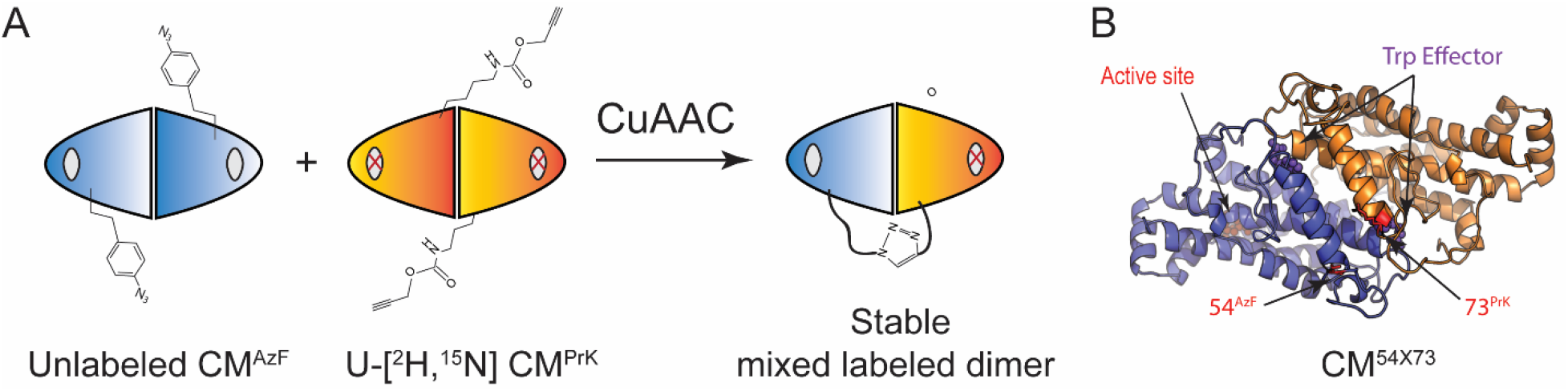
CuAAC approach for generating chorismate mutase mixed labeled dimers. (A) General scheme for stable MLD generation in which Unlabeled AzF-containing protein is mixed with isotopically labeled PrK-containing protein, a Cu^I^ source is added, subunit exchange occurs, and if the two UAAs are close enough, the heterodimer is covalently linked by CuAAC. Note that one subunit has an active site mutation (depicted by the red “X”), which prevents substrate binding in one of the two subunits, thereby facilitating NMR or other structural biology investigation of the elusive singly-bound state. (B) The UAAs used to link CM in this work, 54^AzF^ and 73^PrK^ are shown. The C^β^ to C^β^ distance between these two amino acids is ~ 10 Å in CM x-ray models.

Herein, we overcome these barriers and produce 30 mg of a pure, linked, perdueterated MLD using CuAAC from single liter growths of the two UAA-containing CM dimers. Each of the rudiments used to boost expression of UAA-containing proteins is at least mentioned in the literature, but we show that by carefully manipulating the aaRS plasmid type, the bacterial strain, and the timing and composition of the growth media in combination, one can routinely obtain large yields of protein with different UAA types. Expression is cost effective even for NMR, which requires expensive isotopic labeling, and is notoriously demanding of high concentration samples. By conducting CuAAC in an oygen-free environment, we are able to make the click linkage reaction go to such efficiency, that little to no further purification is required, and the product maintains both the structure and activity of the native enzyme.

## Results and Discussion

### Optimizing expression of AzF and PrK-containing chorismate mutase

Our strategy of using CuAAC to link the two subunits of the CM dimer requires incorporation of an azide-containing UAA in one subunit and an alkyne in the other (Figure 1). UAA incorporation at a single site is achieved through amber codon suppression in concert with engineered tRNA synthetases^20^ with CM overexpression driven by the traditional T7 system in *E. coli*. Given the high mass requirements of NMR spectroscopy samples, optimizing expression yield is a critical element to the success of the approach. For this reason, we chose p-azidophenylalanine (pAzF) as the N3 contributor because the evolved aaRS that incorporates this UAA was shown to be as efficient as the native *E. coli* PheRS^21, 22^. Indeed, expression yields of CM containing pAzF at any one of multiple positions are in range of the wild type when expressed using the pULTRA^23^ plasmid tRNA/aaRS platform (Table 1.). In addition, we observed a significant increase in CM-pAzF protein yield when we used the arabinose inducible pEVOL platform^24^ for expression, presumably due to an increase in the cellular amount of the aaRS. Initially, the high yields lead us to believe that isotopically labeling the N3-containing subunit of the mixed labeled CM dimer would be most economical because the excellent yields of AzF-containing CM would offset the high cost of isotopes. For isotopically labeled applications, we used the approach described by Venditti and co-workers^25^, in which aaRS synthetase expression was pre-induced by arabinose in rich media prior to pelleting the cells and transferring into M9 minimal media supplemented with pAzF and the desired NMR isotopes; yields were outstanding on the order of 50 mg/L (Table 1).

**Table 1.** Expression levels of Chorismate Mutase containing unnatural amino acids

| **E. coli strain** | **aaRS platform** | **UAA position** | **UAA** | **Growth media** | **Yield/L** |
| --- | --- | --- | --- | --- | --- |
| BL21*(DE3) | None | WT enzyme | None | M9<sup>H<sub>2</sub>O</sup> or D<sub>2</sub>O | 50-70 mg |
| BL21*(DE3) | pULTRA/pCNF-RS | 54, 73, 97, 137, 207, 210 | AzF | LB or M9<sup>a</sup> | 20-50 mg |
| BL21*(DE3) | pEVOL/pAzF | 54, 73, 97, 137, 207, 210 | AzF | M9<sup>H<sub>2</sub>Oa</sup> | 50 mg |
| BL21*(DE3) | pULTRA/pylRS | 207 | PrK | LB | 1 mg |
| BL21*(DE3) | pULTRA/pylRS | 207 | PrK | M9<sup>H<sub>2</sub>O</sup> | 1 mg |
| BL21*(DE3) | pEVOL/pylRS | 73, 137, 207 | PrK | M9<sup>H<sub>2</sub>O,b</sup> | 5-10 mg |
| B.95-ΔA | pEVOL/pylRS | 73, 97, 137, 207 | PrK | M9<sup>H<sub>2</sub>O,b</sup> | 5-10 mg |
| B.95-ΔA | pEVOL/pylRS | 73, 97, 137, 207 | PrK | M9<sup>H<sub>2</sub>O,c</sup> | 25-50 mg |
| B.95-ΔA | pEVOL/pylRS | 73 | PrK | M9<sup>D<sub>2</sub>O,c</sup> | 50 mg |
| B.95-ΔA | pEVOL/pylRS | 73 | PrK | M9<sup>+H<sub>2</sub>O,c</sup> | 200 mg<sup>d</sup> |

<sup>a</sup> 1 mM AzF added prior to IPTG induction
<sup>b</sup> Cells grown for two hours in LB + 0.02% arabinose prior to transfer into M9. 1 mM PrK added prior to IPTG induction.
<sup>c</sup> Cells grown overnight in LB + 0.02% arabinose prior to transfer into M9 (or M9+). 10 mM PrK added prior to IPTG induction. Media composition in Supplemental Methods.
<sup>d</sup> Based on yield from a 125 ml culture.

To generate the CuAAC alkyne reactant, we chose to incorporate propargyl-lysine (PrK) into CM due to its relatively low cost and the expected flexibility imparted by numerous bond rotational degrees of freedom. We viewed this plasticity as an advantage, both in terms of satisfying the geometric requirements of CuAAC reactant orientation between the two subunits, and minimizing strain-induced perturbation of the enzyme structural ensemble after linkage. The native pyrrolysine RS (pylRS) from *Methanosarcina barkeri* incorporates the PrK UAA into proteins^26, 27^, and first we used pylRS in the pULTRA platform to express CM with PrK in place of native lysine 207. We obtained 1 mg of 207PrK from either one liter of LB or M9 minimal media (Table 1); this poor yield was accompanied by a large population of truncated product, presumably CM^1-206^ (Figure S1A). To test whether raising the amount of pylRS would increase yield, we subcloned two copies of the gene into the pEVOL UAA expression plasmid backbone^24^, one under the control of the constitutive proK promoter, and the other under the inducible araC promoter (the two aaRS copies is standard in pEVOL). To be sure that the yields were not idiosyncratic to the position of the UAA, we tried multiple substitutions throughout the enzyme (Table 1). Cultures were treated in the same way that gave high yields with the pEVOL-pAzF system above, and while protein yield increased five-fold to ~ 5 mg/L, the majority of the induced protein mass was truncated product (Table 1 and Figure S1B). Truncation at the amber codon is often seen in UAA protein expression, likely due to effective competition of release factor (RF1) with the charged amber tRNA^28^. We therefore utilized a strain (B.95-ΔA), which lacks RF1 to alleviate the truncation phenomenon, and replaces UAG codons in select genes with other stop triplets to maintain fitness^29^. Unfortunately, while there is no truncated product induced in this strain, yields are still on par with the classic BL21*(DE3) cells (Table 1 and Figure S1B), a result that is also independent of UAA position. It was previously shown that increasing the [PrK] in the expression media from 1 mM to 10 mM increases expression^26^. To go along with higher aaRS substrate, we hypothesized that extending the time of growth in rich media with arabinose from two hours to overnight would increase the concentration of the aaRS enzyme itself. These two changes, coupled with the B.95-ΔA strain, were all necessary to obtain yields between 25 and 50 mg/l, which is on par with the wild-type and AzF UAA enzymes (Table 1 and Figure S1C). This high yield was maintained when cells were grown in M9 with 99.8% D_2_O, which is critical for NMR studies of this 60 kDa enzyme (Table 1) and will enable CuAAC NMR applications to larger systems. Lastly, perdeuterated growths of PrK-containing enzymes can be quite costly owing to the D_2_O at $400/L and the UAA at $250/L. Clore and coworkers developed a minimal media, called M9+, in which both the levels of glucose and the buffering capacity are raised, allowing cells to grow to 10-fold higher density than canonical M9^30^. We adapted our overnight pre-induction in LB-arabinose approach to M9+ (See Methods) and obtained 200 mg/L, which equates to a 4-8-fold cost savings on a per mg of expressed protein basis (Table 1). Each of the principles applied here were foreshadowed separately in the literature, but we show the importance of employing them in unison. Collectively, the insights described above can be applied to NMR and other structural biology applications where cost effective production of UAA-containing proteins was previously a barrier.

The CuAAC reaction is notoriously difficult in air and aqueous solution due to the damaging effects of the ROS cascade on macromolecules. Thus, it was important for us to find conditions that are compatible with both efficient CuAAC linkage and maintenance of enzyme integrity post linkage. To test the two CM CuAAC linkage partners independently, we reacted either CM^73PrK^ or CM^54AzF^ with a 5 kDa PEGylated azide or alkyne respectively (Figure S2A). We observed efficient CuAAC with CM^73PrK^ and mPEG-Azide in air using CuSO4 as the copper source and ascorbic acid as the reductant (Figure S2B). Unfortunately, despite taking precautions to limit enzyme damage by ROS, including degassing the reagents, floating Argon on the reaction, using excess Cu^I^ ligand (BTTAA)^31^, and adding aminoguanidine to the reaction^32^, CM lost nearly half of its catalytic activity under these conditions (Figure S2C). In order to avoid the ROS-generating reducing agent, we moved to a Cu^1^ source in the form of [(CH_3_CN)_4_Cu]PF_6_, and while exposing CM to this reagent in air did not affect activity (Figure S2C), the CuAAC efficiency was severely impaired (Figure S2D), likely due to rapid loss of catalyst upon oxidation of Cu^I^ to Cu^II^. We therefore moved to anaerobic conditions^19, 33, 34^ and performed the reaction using the Cu^I^ source, [(CH_3_CN)_4_Cu]PF_6_, in a glove box under N2 (See Methods). Under these conditions, we observed both efficient CuAAC reaction and preservation of enzyme activity (Figure S2C and S2D). It also became clear from these test reactions that the CM^73PrK^ enzyme was reproducibly more reactive than CM^54AzF^, yielding CuAAC efficiencies of ~90% and ~60%, respectively (Figure S2E). We found that using the AzF-containing protein immediately after purification (within days of lysing cells) maximized the reactivity, but CM^PrK^ was always more reactive than CM^AzF^, even after the PrK protein was more than a month old. We were careful not to expose CM^AzF^ to reducing agent, and mass spec confirmed that azide reduction by this mechanism was not the culprit. Nonetheless, the observation steered our MLD linkage strategy towards isotopically labelling the PrK subunit so any unlinked AzF-containing CM that we could not purify away from linked dimers would be silent in NMR experiments.

To generate a MLD for NMR spectroscopy, we used unlabeled 54^AzF^ and U-[^2^H,^15^N] 73^PrK^ (natively Asn and Arg, respectively). The C^β^-C^β^ distance between these two amino acids is ~10 Å in x-ray models (Figure 1B), which should be compatible with subunit linkage. Immediately prior to preparative scale CuAAC, we ran small scale PEG-reactions and found that 54^AzF^ and U-[^2^H,^15^N] 73^PrK^ were ~70% and ~90% reactive, respectively (Figure 2). In addition, a titration of the relative concentrations of linkage partners in CuAAC reactions showed that a 54^AzF^:73^PrK^ ratio of 1.5:1 yielded maximal linked product (Figure 2). We should note that product peaked after 30 minutes in the presence of Cu^I^, indicating rapid mixing of CM subunits. The conclusion that mixing induced pre-organization greatly accelerates and is necessary for CuAAC under our conditions, is supported by our observation that while CuAAC is successful on the dimer at 25 μM total protein concentration, one of the click partners needed to be at millimolar concentration in the bi-molecular PEG linkage reactions (Figure S2B). The preparative reaction contained 32 mg of 54^AzF^ and 21 mg of 73^PrK^ mixed in an anaerobic glovebox (see Methods) giving outstanding linkage efficiency (Figure 2A). Centrifugation of the product lead to an enrichment of linked product due to selective precipitation of the *unlinked* azide (data not shown). We were alerted to the potential utility of this phenomenon by witnessing precipitation of CM^AzF^ in the presence of Cu^I^ without its CuAAC PrK partner, while CM^AzF^ remained clear in the presence of Cu^I^, as long as there was excess alkyne. We obtained a yield of 33 mg product (62% of input mass), of which nearly all was linked (Figure 2A). This is close to theoretical maximum (66% of the total input mass) based on the 73^PrK^ limiting reagent, consistent with the near complete reactivity of this linkage partner (Figure 2A). Finally, although NMR spectroscopy showed no evidence of detectible ^15^N unlinked 73PrK homodimer (see below), we were able to achieve additional purification either by adding an excess of His-tagged CM to the linkage reaction, after which unlinked material dimerized with tagged subunits and bound to Ni^2+^resin (Figure 2A). Although it was not necessary in this labeled preparation, we are able to purify away unlinked material by size exclusion chromatography in sub-denaturing urea if necessary (Figure S3). The linked material is a homogeneous dimer in solution, in other words, there is no trans-dimerization as shown by indistinguishable SEC traces of concentrated linked and unlinked CM under native conditions (Figure S4). Lastly, the mixed dimer retains all the catalytic features of the wild-type unlinked enzyme. In the absence of effector, the V_max_, *K_m_*, and Hill coefficients are indistinguishable (Figure 2B and Table S1). This is an important detail as it shows linkage does not perturb the classical homoallostery we intend to study with this linked MLD. The linked species is also activated by tryptophan and inhibited by tyrosine, though there are subtle differences between linked and unlinked CM with respect to heteroallostery (Figure 2B and Table S1). This is not surprising given the location of the linkage in the effector binding domain (Figure 1B).

**Figure 2.**
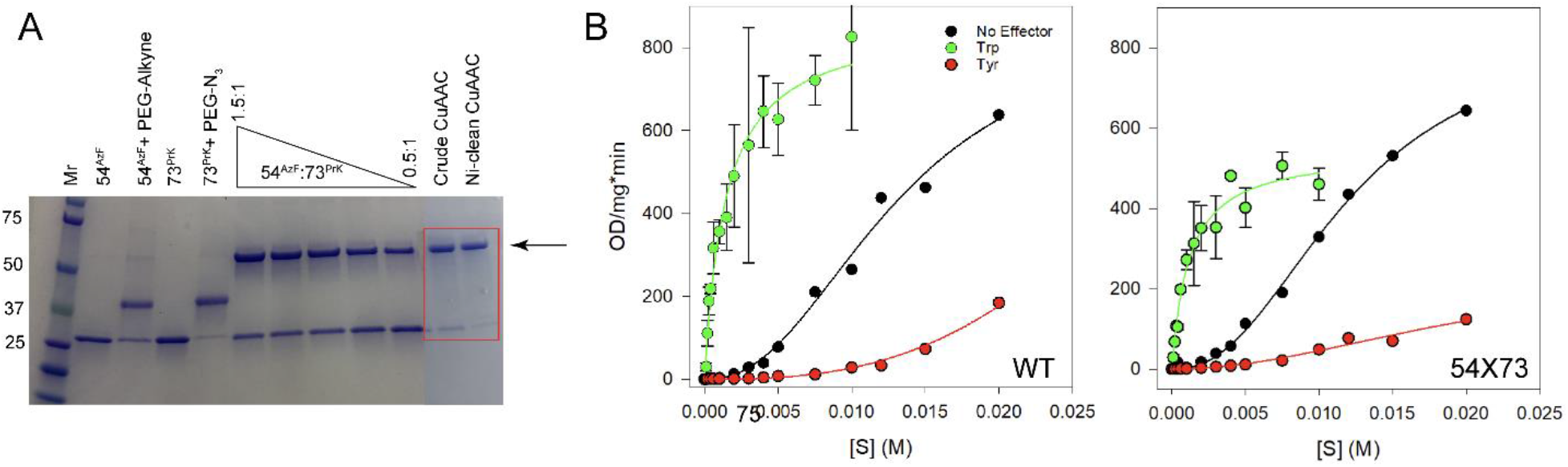
CuAAC and enzymatic activity of linked CM. (A) Demonstration of using CuAAC to link subunits of 54^AzF^ and 73^PrK^ CM. First four lanes show the reactivity of the individual subunits by using PEG click reagents, followed by small scale titration to establish the optimal ratio of the two subunits. The arrow shows the position of the linked product. The penultimate lane is a crude sample from a preparative 53 mg reaction, and the final lane shows modest purification after using His-tagged CM to fish out unlinked CM (see Methods). (B) Activity of unlinked wild-type CM in the left panel, and linked 54X73 CM in the right shows that linkage has no effect on activity of a the apo enzyme, along with subtle effects on the activity in the presence of Trp and Tyr effectors.

Consistent with activity, NMR spectroscopy shows that neither exposure to copper, nor mutation, nor linkage itself significantly affects the structure of CM. In what follows, we denote the linked mixed labeled dimer between 54^AzF^ and 73^PrK^, 54X73*, where the asterisk indicates that only the 73^PrK^ subunit is perdeuterated and labeled with ^15^N. This approach of isotopically labeling, and therefore observing only one subunit at a time, is a prerequisite of our MLD approach for studying allosteric binding intermediates. First, we were able to concentrate 54X73* to 500 μM dimer and collect a TROSY ^1^H-^15^N HSQC spectrum, which is very similar to spectra of symmetrical 73^PrK^ (Figure 3A) and wild-type (Figure 3B) homodimers. However, we observe less than 200 out of the possible 242 non-proline resonance in *wild-type* spectra, and these missing peaks largely map to the effector binding domain (EBD), which is the location of the 54 to 73 linkage (Figure S5). Thus, in principle, we could miss an effect of linkage on EBD structure because these peaks are missing. To test this possibility, we synthesized the CM transition state inhibitor^35, 36^, and added it and the tryptophan activator to wild-type or 54X73* CM and acquired HSQC spectra of this so-called “super-R” state^10^. Contrary to the apo, tryptophan, or tyrosine bound states, the super-R state gives rise to nearly all possible backbone resonances in HSQC spectra, and comparisons of these fingerprints shows that neither mutation nor linkage significantly affects the structure of the EBD (Figure 3C and 3D). Lastly, our strategy of labeling the PrK-containing protomer of the MLD was predicated on the fact this amino acid was consistently nearly 100% reacted in test CuAAC reactions (Figure 2A and S2E), thereby placing less stringency on the need to separate out linked material because the unreacted, unlabeled AzF partner would be undetectable. While our preparations of linked MLD are highly pure, we do see a small amount of unlinked material on Coomassie stained gels (Figure 2A). However, careful analysis of 54X73* HSQCs shows no intensity attributable to the 73* homodimer, suggesting the small amount of unlinked CM is silent in NMR experiments as per the design (Figure 3C, inset). The advances described in this work will certainly be useful in elucidating allosteric mechanisms in allosteric dimeric and higher order oligomer systems. It is our hope that the ability to obtain high yields of intact and homogeneously cross-linked protein will also enable structural and dynamic insights into the function of otherwise intractable systems such as transient protein-protein interactions and large molecular assemblies^37^.

**Figure 3.**
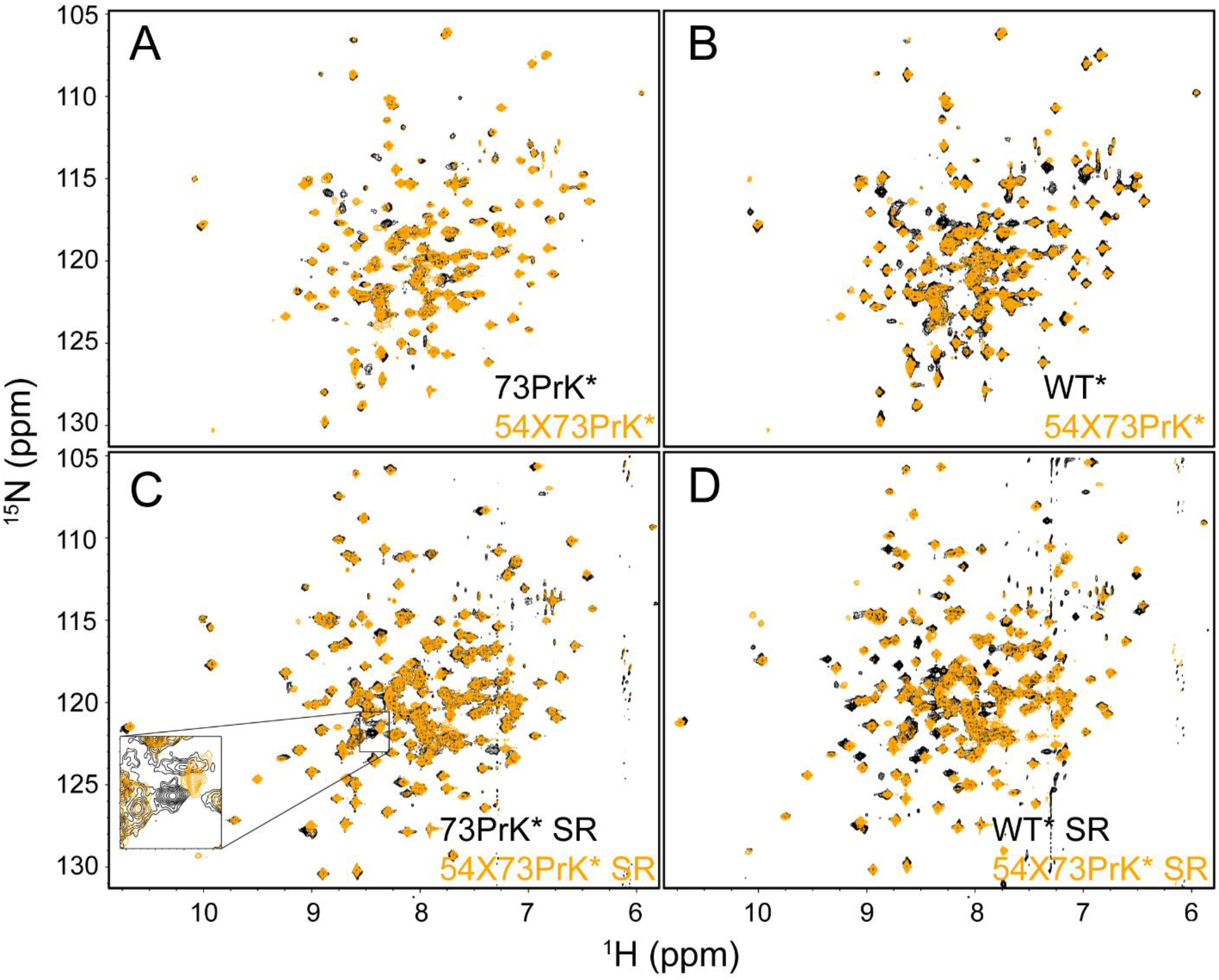
NMR spectroscopy of the 54X73* linked mixed labeled dimer. (A) Comparison of TROSY ^1^H-^15^N HSQCs of apo 54X73* with the 73PrK* homodimer. (B) Comparison of apo 54X73PrK* MLD with the free wild-type enzyme. Collectively panels A and B shows that neither the mutations nor the linkage significantly affects the structure or dynamics of CM. (C) Comparison of the 54X73 * and 73PrK* super-R (SR, tryptophan plus transition-state inhibitor-bound ternary complex). The inset in panel C shows that there is no detectable unlinked 73PrK* homodimer in our linked MLD preparation as this contaminant would represent orange intensity overlaying with the central black resonance. (D) Comparison of the WT and 54X73 * MLD super-R spectra. Note that the MLD is not “back exchanged” (^2^H to ^1^H of highly protected amides), which explains the ~10 back resonances without an orange counterpart.

## Methods

### Expression and purification of CM

TEV cleavable His-tagged wild-type CM in pET21a was transformed into BL21*(DE3) cells, grown to OD_600_ ≅ 0.8 in various liquid media types at 37 °C, IPTG was added to 0.75 mM, and induced for 4 hours. Cells were collected in Ni^2+^ A buffer (20 mM NaPO_4_, 500 mM NaCl, 10 mM BME 0.02% NaN3, pH 7.4), 10 mg hen egg white lysozyme was added, and cells were frozen at −20 °C until further use. Cells were lysed using sonication, and the cleared supernatant was applied to a 5 ml Ni^2+^ column (Genesee or Cytivia) and eluted with a 15 column volume gradient from 0-100% Ni^2+^ B buffer (Buffer A plus 250 mM imidazole). CM-containing fractions were pooled, roughly 0.3 mg of TEV protease was added, and the cleavage was allowed to proceed overnight while dialyzing against Buffer A at room temperature. Cleaved CM was added to a Ni^2+^ column, the flow through was collected, concentrated, and then applied to an S200 size exclusion chromatography column equilibrated with NMR buffer (25 mM NaPO_4_, 150 mM NaCl, 1 mM EDTA, 0.01% NaN_3_, 1 mM DTT, pH 7.5).

For growths involving UAAs, the CM plasmid above containing an amber mutation was co-transformed with an aaRS/tRNA plasmid (either pULTRA or pEVOL) into BL21*(DE3) or B.95-ΔA cells. Growths using the pULTRA platform proceeded similarly to the wild type because the aaRS is under the control of a constitutive promotor in this system. To express AzF-containing CM in the pEVOL system, a 150 ml culture of LB was inoculated with 5 ml of an overnight culture, and grown at 37 °C. When the cells reached an OD600 of 0.4, arabinose was added to 0.02%, and the cells were grown for an additional two hours prior to collection by centrifugation at 170xg for one hour. These cells were then added to one liter of M9 minimal media such that the OD was 0.4 (cells from the entire 150 ml overnight culture were not always necessary to reach 0.4), arabinose was added to 0.02% and AzF was added to 1 mM, and the cells were allowed to grow until the OD reached 0.8, at which time IPTG was added and induction proceeded for four hours as described above. Expression of PrK-containing CM in minimal media was conducted similarly except the initial 150 ml grown in LB-0.02% arabinose proceeded overnight rather than two hours, the unnatural amino acid was added to a [final] of 10 mM rather than 1 mM, and induction time was extended to 12 hours. Expression in 99.8% D_2_O proceeded similarly with the following modifications: the M9 media was supplemented with 1 g/L ^2^H, ^15^N celtone powder (Cambridge Isotope Labs), and 1X MEM Vitamins (Fisher). In addition, cells grown in LB-arabinose were washed twice with M9 salts in D_2_O to remove as much rich media as possible prior to transfer into 99.8% M9. Lastly, as a cost saving measure, we demonstrated the ability to get high yield of UAA proteins in a higher cell density, lower culture volume media called M9+. The composition of M9+ was as described by Clore^30^ and as is the case with UAA growths in classic M9 above, we generated the majority of the cell mass in LB-arabinose prior to transfer into M9+ to avoid suppression of transcription from the pBAD promoter by glucose. Cells were washed with M9 salts, added to M9+ such that the OD was 4.0, arabinose and the appropriate UAA were added, and cells were grown to an OD of 8.0 prior to induction. Collection and purification of UAA containing CM was as above with the wild type except reducing agent was omitted from all steps involving AzF protein in order to avoid reducing the azide, and NaN3 and reducing agent was omitted from all final buffers so as not to interfere with downstream CuAAC applications.

### CuAAC

All CuACC reactions were conducted in 25 mM HEPES, 150 mM NaCl, pH 7.5, without reducing agent. Control reactions were conducted with either mPEG-azide (MW 5k) or mPEG alkyne-alkyne (MW 5k) from Creative PEG Works. PEG CuAAC reactions with CuSO_4_ contained 10 μM CM^AzF^ or CM^PrK^ dimers, 5 mM aminoguanidine, 0.5 mM CuSO_4_, 2.5 mM BTTAA, and 5 mM ascorbic acid, added in that order. Alternatively, reactions with Cu^I^, substituted [(CH_3_CN)_4_Cu]PF_6_ for CuSO_4_ and omitted the ascorbic acid. [(CH_3_CN)_4_Cu]PF_6_ was prepared in acetonitrile, mixed with BTTAA prepared in water, lyophilized, and then re-suspended in degassed buffer in air or in a glovebox. A titration showed the concentration of the PEG reagents for optimal linkage was 5 mM (Figure S2B). Reactions were performed either in air or in a glovebox (Cleatech) under N_2_, and were allowed to proceed for 30 minutes prior to adding EDTA to 10 mM. Intra dimer CM linkage reactions used 2-fold lower copper:BTTA (0.25:1.25 mM) and both protein dimers (AzF and PrK CuAAC partners) were on the order of 10 uM. The preparative mixed labeled dimer reaction was 35 ml in which the 54^AzF^ and 73^PrK^ concentrations were 15 and 10 μM dimer respectively. EDTA (10 mM) was added to the reaction in the glovebox and to remove the copper-EDTA chelate, the material was immediately buffer exchanged into NMR buffer.

### Separation of linked MLD from unlinked

To clean up the MLD sample, we added 10-fold excess His-tagged CM to our crude MLD preparation, and allowed it to mix for one hour at room temperature in Ni-buffer A. The scale of this separation was 15 mg MLD, of which, we estimated 10% unlinked, therefore 15 mg of His-CM was added. The mixture was then applied to a 5-ml Ni^2+^ column (Cytivia), and the enriched MLD, which lacked a His tag, was in the flow through (Figure 2A), while unlinked subunits paired with His-tagged subunits and adsorbed to the column. We explored an alternative purification method of adding 4 M urea to the crude MLD, and running it over an S200 size exclusion column equilibrated with NMR buffer plus 4M urea. This was also effective (Figure S3), but urea exposure resulted in ~25% loss of the material.

### CM Activity Assay

The substrate, chorismic acid (CA) from Sigma was suspended in D_2_O, and its concentration was determined by the NMR-based PULCON experiment^38^. CA was then lyophilized and re-suspended in NMR buffer, and base was added to ensure accurate and constant pH in experiments with different CA concentrations. Conversion of CA to prephrenate is accompanied by maximal absorbance change at 274 nm (Figure S6A). However high concentrations of CA in Michaelis-Menten type analysis will result in A_274_ beyond the linear range of our spectrophotometer. To get around this, we measured kinetics at a non-maximal wavelength. In other words, we targeted the shoulder and measured kinetics at 310 nm. To ensure this is justified, we measured the extinction coefficient for the substrate to product conversion at a range of wavelengths, and found that the extinction coefficients were precisely determined and constant over a range of substrate concentrations (Figure S6).

To monitor CM activity during simulated CuAAC conditions, we exposed the wild-type enzyme to the reaction conditions described in methods above, an aliquot was removed at the specified time, EDTA was added to 5 mM, the aliquot was centrifuged, and activity was measured in NMR buffer under the following conditions: 40 nM CM, 1 mM CA, and 10 μM Trptophan. A310 was monitored for one minute, and the initial velocities were determined from the first 100 ms. V/V_o_ was then plotted as a function of time, where V is the initial velocity at time *t* after addition of copper, and Vo is the initial velocity in the absence of copper.

Full Michaelis-Menton analysis was performed on the 54X73* MLD and compared with the wild-type. All assays were performed in NMR buffer at 22 °C and velocities were measured at 310 nm as described above. Due to the dramatically different velocities in the absence and presence of effector molecules, we modulated the enzyme concentration to bring the kinetics into a favorable window for measurement. The apo enzyme was at 4 E^-7^M, the Trp-bound enzyme was at 1.5 E^-8^ M CM and 40 μM Trp, and the Tyr-bound enzyme was at 8 E^-7^ M CM and 100 μM Tyr. Velocities were normalized to the mass of enzyme and plotted as velocity per mg CM in Figure 2B.

### NMR Spectroscopy

500 μM (dimer) U-[^2^H, ^15^N] WT or 54X73* samples were prepared in NMR buffer supplemented with 5% by volume D_2_O. TROSY-HSQC spectra of either were acquired at 25 °C on an 600 MHz Bruker Avance III spectrometer equipped with a TCI 5 mm cryoprobe using the trosyf3gpphsi19.2 Bruker pulse program with 16 scans per fid, a recycle delay of one second, and (1200, 110) complex points corresponding to (100 ms, 64 ms) acquisition times in t2 and t1, respectively. Tryptophan was added to 7.4 mM and two molar equivalents of the transition state inhibitor were added prior to acquisition of HSQC spectra of the super-R state using the same parameters as above.

