## Supplementary Info for "CuAAC stabilization of an NMR mixed labeled dimer"

### Supporting Information

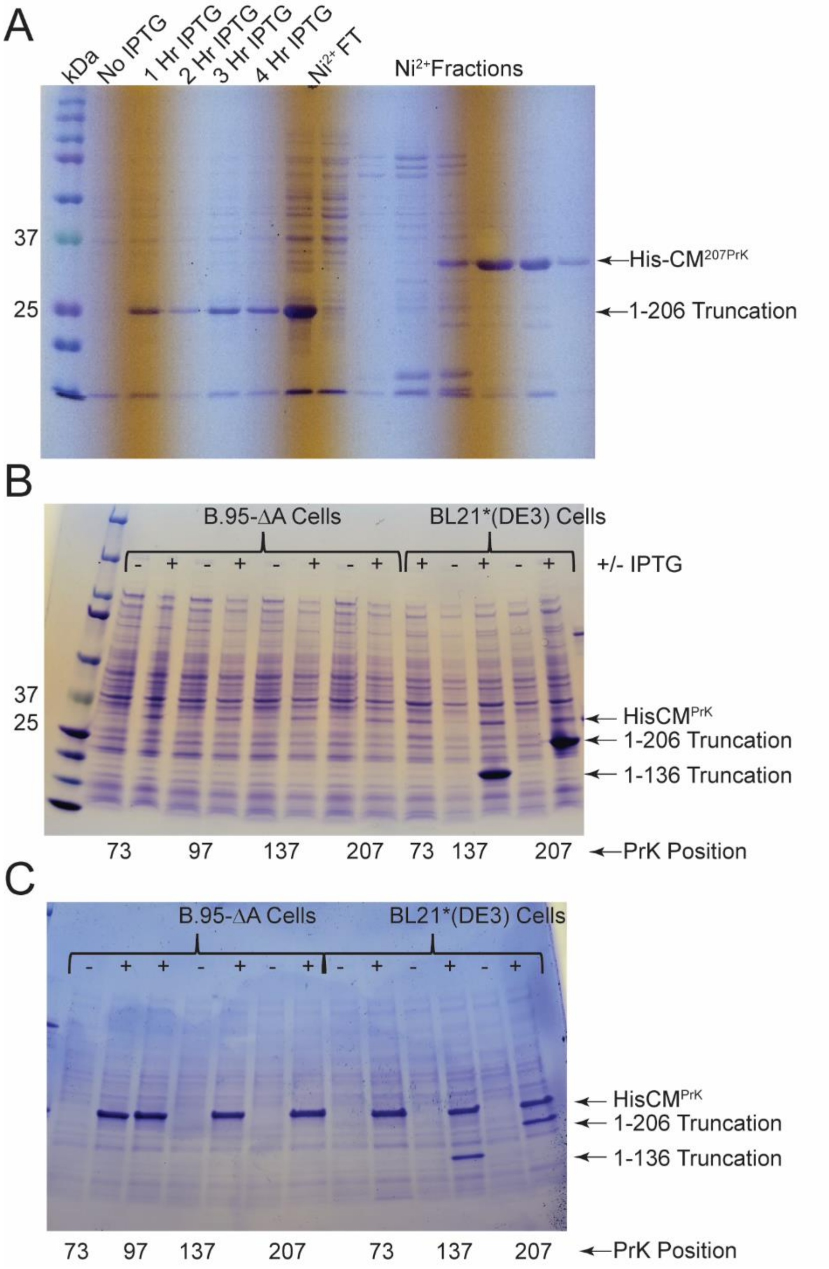


*Figure S1 Parameters affecting expression of PrK-containing CM.* (A) Purification of $CM^{207PrK}$ expressed from BL21*(DE3) cells transformed with the tRNA/pylRS containing pULTRA plasmid in a 1 liter LB culture. Lanes show a time course of expression as well as the $Ni^{2+}$ column purification. The yield from this growth was 1 mg/L (See Table 1 from main manuscript). (B) Effects of using the pEVOL platform to express the tRNA/pylRS pair and the impact of the RF1 knockout strain, B.95-ΔA, on PrK-containing CM. Note that the pEVOL platform enhances expression (compare induced bands in panels B vs A) relative to pULTRA, and the while the B.95-ΔA strain ameliorates the truncation problem, pEVOL was sufficient to increase yield to 5-10 mg/L (See Table 1). (C) Effect of increasing the arabinose pre-induction duration from 2 hours to overnight and increasing the [PrK] in cultures from 1 mM to 10 mM. Intensity of expression bands show that both of these changes resulted in a further increase in yield to 25-50 mg $CM^{PrK}$ per liter (See Table 1). Using the B.95-ΔA strain eliminates the truncation problem and gives yields on the higher end of the 25-50 mg/L range.

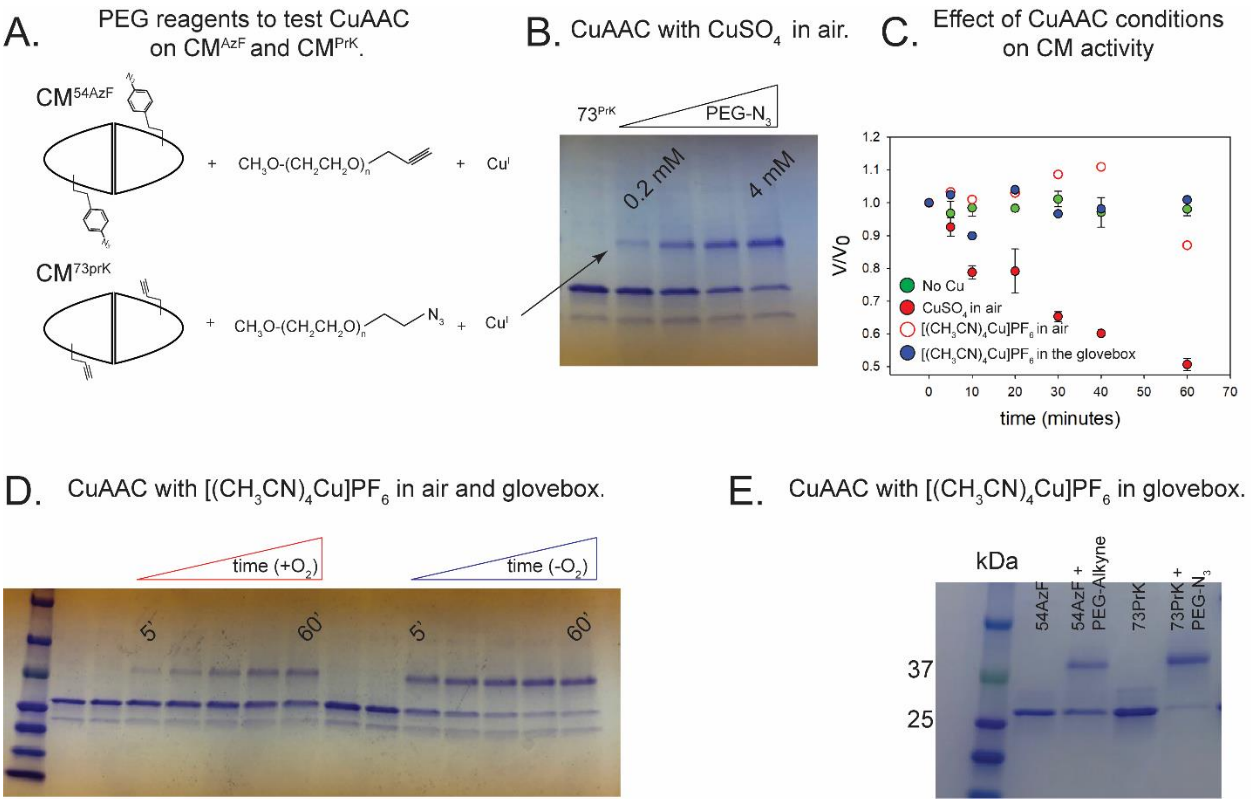


*Figure S2 Optimizing CuAAC for linkage efficiency and maintaining function.* A) mPEG-Alkine (5 kDa) and mPEG-$N_3$ reagents used to test CuAAC reactivity of $CM^{AzF}$ and $CM^{PrK}$, respectively. (B) Optimizing PEG reagent concentration required for efficient CuAAC using $CuSO_4$ and ascorbate. Reaction conditions were: 10 μM CM, 0.5 mM $CuSO_4$, 2.5 mM BTTAA, 5 mM aminoguanidine, 5 mM ascorbate, 150 mM NaCl, 25 mM HEPES, pH 7.5. (C) Wild type CM activity after enzyme exposure to various CuAAC reaction conditions. EDTA was added after the prescribed time, activity was measured, and compared to the velocity at time 0. (D) Comparison of CuAAC efficiency of $CM73^{PrK}$ with PEG-$N_3$ using the $[(CH_3CN)_4Cu]PF_6$ catalysis out (red) and in (blue) of an anaerobic chamber. Note the increased efficiency in the glovebox due to maintenance of the proper +1 oxidation state of the catalyst. The first two lanes in each series are $CM^{73PrK}$ with and without $Cu^I$ in the absence of PEG reagent. (E) $CM^{54AzF}$ (lane 1) was mixed with mPEG-Alkyne (5 kDa) and $[(CH_3CN)_4Cu]PF_6$ in an anaerobic glovebox, and reacted until the amount of product reached a maximum (lane 2). A similar experiment was performed with $CM^{73PrK}$ (lane 3) in a CuAAC reaction between $CM^{73PrK}$ and mPEG-Azide (5 kDa). Note that the $CM^{54AzF}$ reaction went to about 50% and the $CM^{73PrK}$ went to about 90% completion. Although reactivity of $CM^{AzF}$ could exceed 50% if we used this protein within days of purification, the difference in reactivity between $CM^{N3}$ and $CM^{PrK}$ was consistent in multiple protein preparations with UAAs at multiple positions.

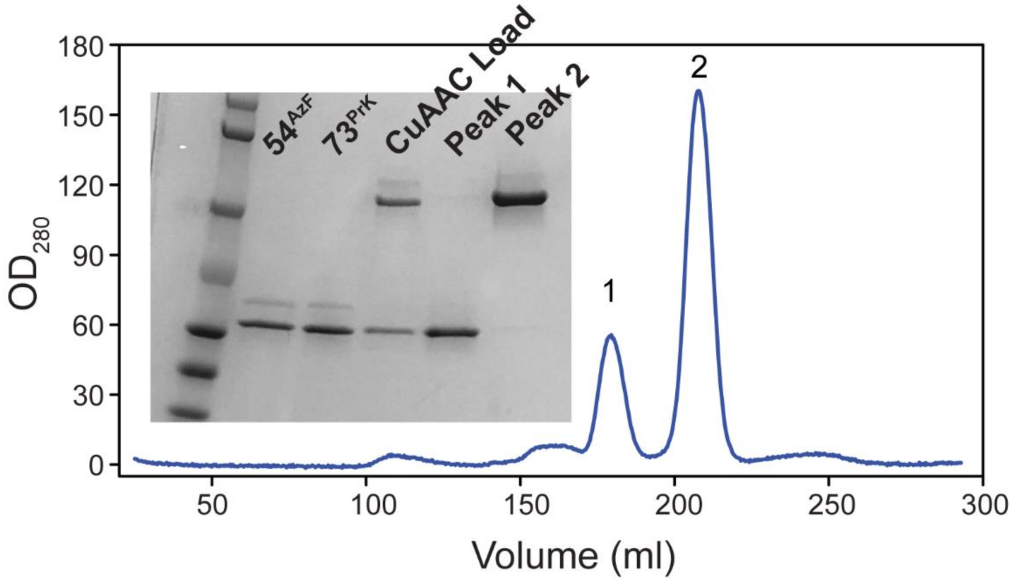


**Figure S3.** Separation of linked 54X73 from unlinked $54^{AzF}$ and $73^{PrK}$ using size exclusion chromatography in 4 M urea. 15 mg of crude CuAAC reaction was loaded onto an S200 column, peaks 1 and 2 were individually pooled and run on an SDS gel (inset) showing near perfect separation.

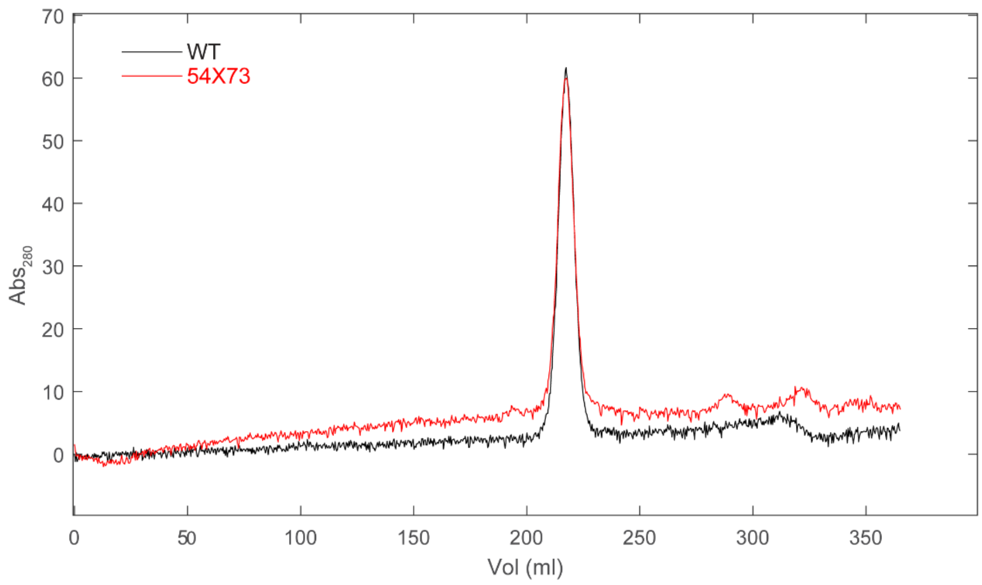


**Figure S4.** Linked CM is a dimer in solution. WT and 54X73 (80 μM dimer) were run separately over a S200 size exclusion column, and the linked material elutes at the exact same position as the wild type. This indicates that there is no trans-dimerization or higher order oligmerization between linked dimers.

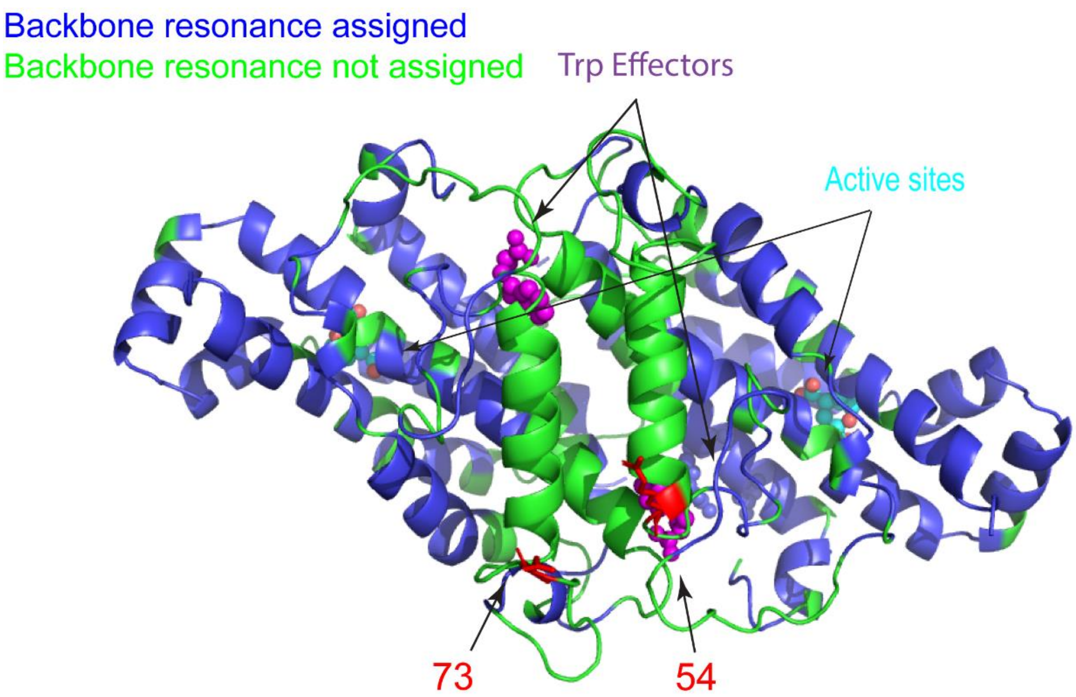


**Figure S5.** The backbone of most CM states is not assigned in the effector binding domain (EBD). Assigned (blue) and unassigned (green) amide resonances of the CM-Trp are shown on the x-ray model. Note that the position of the 54 to 73 linkage described in this work is in this domain. The unassigned peaks are missing, so it is possible we could be blind to the effect of linkage on this domain. Nearly all resonances are visible in the Super-R state (Trp and Transition-state-inhibitor bound). Overlay of wild type and 54X73* Super-R spectra show that the inhibitor binds to the linked dimer, and the structure of the entire protein is preserved (Figure 3D, main text).

Table S1. Enzymatic activity of unlinked and linked CM[a].

| | Apo | | | Trp | | Tyr | | |
|---|---|---|---|---|---|---|---|---|
| | $K_m$ (mM) | $V_{max}$ (OD/mg*min) | Hill | $K_m$ | $V_{max}$ | $K_m$ | $V_{max}$ | Hill |
| WT | 13.2 | 868 | 2.30 | 1.4 | 870 | NF[b] | | |
| 54X73 | 12.0 | 845 | 2.33 | 1.2 | 546 | 22 | 267 | 1.99 |

[a] V vs [S] plots in Figure 2.

[b] Fit not reliable due to undefined $V_{max}$.

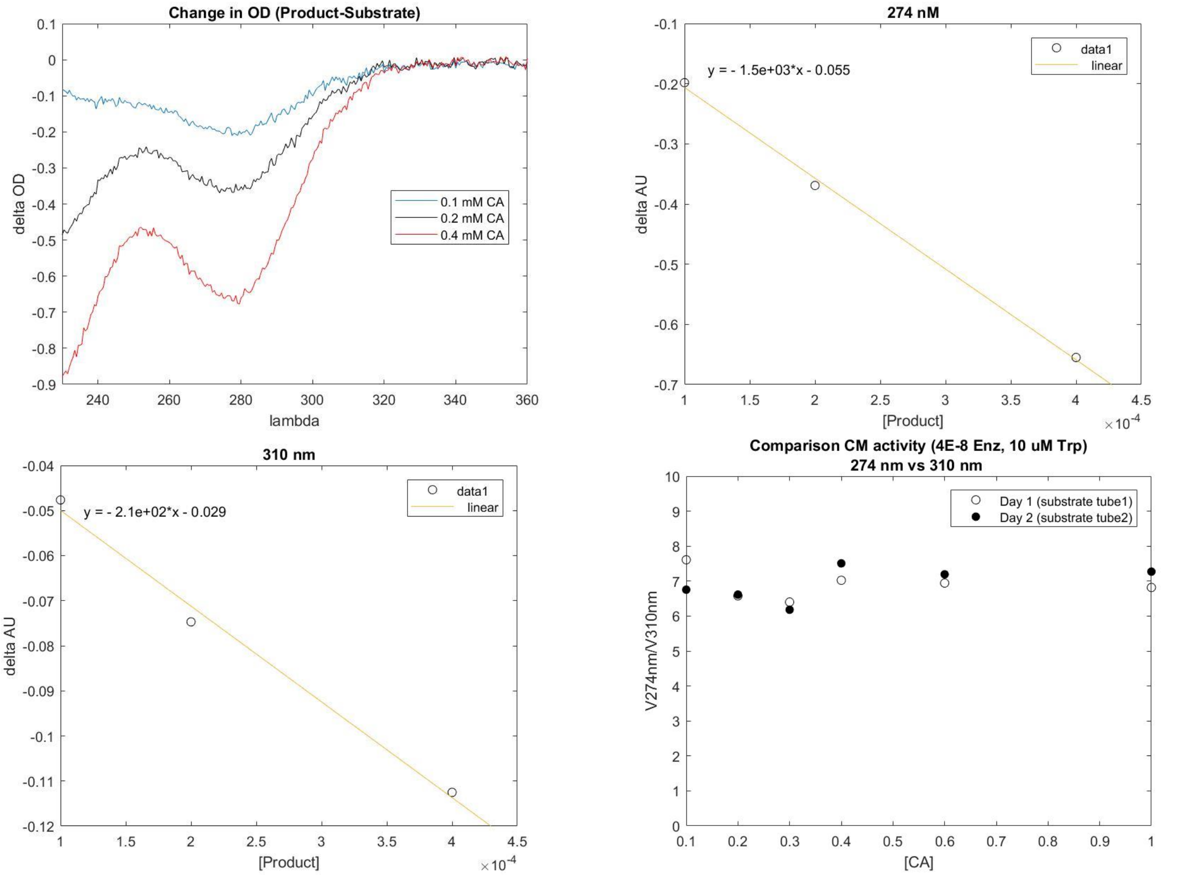


Figure S6. Tests showing robustness of non $\lambda_{max}$ monitoring of CM reaction velocity. Top left: CM catalyzed change in absorbance as a function of wavelength at multiple chorismic acid concentrations. These data can be used to determine the extinction coefficient for the chorismic acid to prephenate conversion at any wavelength as is shown in the 274 nm and 310 nm panels above. The consitent ratio of slopes measured at 274 and 310 nm over the range of CA concentrations further supports conducting the activity assay at wavelength on the shoulder of the peak.
